# Physical forces drive *C. elegans* embryonic deformation

**DOI:** 10.1101/2024.03.19.585678

**Authors:** Ting Wang, Martine Ben Amar

## Abstract

The abnormal development of embryos is closely linked to abnormal cell division and elongation, but the underlying mechanism remains to be elucidated. The embryonic development of *C elegans* embryo is different because it occurs without cell proliferation or cell rearrangement. Here, we focus on a spectacular 4-fold elongation that is achieved approximately 3 hours before the egg shell hatches and results from active filament networks. The body shape is represented by an inhomogeneous cylinder, which allows us to assess the active stresses induced by the actomyosin network located in the cortex and the muscles in ventral position near the epidermis. By considering the specific embryo configuration, we can quantitatively obtain the contractile forces induced by actomyosin filaments and muscles for a bending torsion event with defined curvature. We find that the active stress induced by actomyosin molecular motors or muscles increases with elongation and bending curvature, while also varying with radius. Both elongation and torsional deformation contribute to increased moment magnitudes that explain the dynamics of the embryo in the egg. Our results highlight the complex interplay between biomechanical factors in modulating embryonic deformation.

## 1 Introduction

In nature, the forms of organisms vary widely, but in the early stages of embryonic development, different organisms show a high degree of similarity, revealing the similar origin of biological evolution. One of the typical developmental phenomena is the bending and stretching of embryonic structures.

This phenomenon occurs in the early embryonic development of many organisms, such as the elongation of the mouse embryonic primitive streak (Kyprianou et al., 2020), the elongation of the midgut of the Drosophila embryo (Keller, 2006; Mitchell et al., 2022), and the elongation of the body axis of *C. elegans* (Ding et al., 2004; Lardennois et al., 2019; Priess and Hirsh, 1986). The abnormal development of the fetus is also closely related to the abnormal elongation and division of the embryo during development, but the underlying mechanism remains to be explored. Understanding the deformation behaviour of biological embryonic development and exploring the underlying mechanism of its abnormal development are crucial to new strategies for the treatment of related diseases. Previous work has focused on genes and other biomolecules that shape bodies, while mechanical forces have received much less attention (Dance, 2021). Physical forces play a central role in tissue morphogenesis (Ben Amar et al., 2019; Gilmour et al., 2017; Kuhl, 2016; Molnar and Labouesse, 2021; Tallinen et al., 2016) and animal development (Fei et al., 2020; Vuong-Brender et al., 2017; Valet et al., 2022; Zhang et al., 2011), including active forces exerted by molecular motors and muscle activity, and passive mechanical resistance forces due to tissue elasticity (Ackermann et al., 2022). Understanding how physical forces influence morphogenesis and embryonic development remains to be elucidated.

In recent decades, morphoelastic events driven by active filaments have been proposed to be responsible for the shape evolution of embryos. Biological model systems such as Drosophila or *C. elegans* have brought together a large community of biologists working on the same species to elucidate all aspects of morphogenesis. From a theoretical point of view, the description of biological passive tissues is now well established and biomechanics offers a wide range of models able to treat all kinds of materials, inhomogeneous and/or anisotropic. The complexity of the modeling, given by a constitutive relationship between elastic stress and elastic strain or by a free energy given in terms of invariants, has been tested in detail, sometimes confronted with real tissues. However, at embryonic stages, experimental tests between real tissues and models *in vivo* are extremely delicate (Vuong-Brender et al., 2017), if not impossible, and the validity of the approach is judged by its ability to explain and perhaps quantify biological features. Drosophila and *Caenorhabditis elegans* are good examples of this, as they undergo a series of transformations in a short period of time. These shape changes are driven either by growth or by active filaments such as actomyosins and muscles. Growth is a slow process compared to any dynamical time scale, especially compared to energy dissipation, and is often treated as a plastic transformation of bodies, a topic well established in biomechanics (Dervaux and Ben Amar, 2008; Goriely and Ben Amar, 2005; Rodriguez et al., 1994). As far as active filaments are concerned, two different formalisms coexist without any real study of their equivalence in the general case: either the loads developed by the muscles are represented by an active stress or by an active strain. In the first case, the equilibrium equations are modified, while in the second case it is the elastic strain that is modified in the same way as the volumetric growth. Active strain or active stress give different results even in a uniaxial shear (Giantesio et al., 2019) and the results also depend on the constitutive equations of the material. The resolution of the ambiguity between the two models also seems to depend on the material studied (squeletal (Klotz et al., 2021) or smooth muscle (De Vita et al., 2017)), but little is known about the muscle constitution at the embryonic stage.

On the other hand, to return to the objective of this study, embryonic transformations are not so easy to describe mathematically: the *C. elegans* embryo is confined in an egg before hatching and its elongation occurs with bending, rotation and torsion of its body after a first period of elongation induced by the actin filament. This first period has been studied recently and its residual strain is treated here by a pre-strain formalism. We focus on the second period, which is experimentally characterized by a rather strong agitation of the worm inside the egg, and we modify our strategy by considering active stresses due to actomyosin filaments and muscles. Even if we limit the geometrical description of the embryo to its body shape, *i*.*e*. to a cylindrical shell representing the epidermis, this is composed of different cellular bands with different mechanical and biological properties. In particular, actin filaments have a different geometry and molecular motors seem to be more concentrated in the seam than in the DV cells. The complexity of the elastic formalism is therefore correlated to the geometric deformations of the body shape. This also limits any resolution by computational software.

This paper quantitatively investigates the physical forces on the bending and torsion of the *C. elegans* embryo and is organised as follows. In Sec.2, the last stage of the *C. elegans* embryo just before hatching is explained. In Sec. 3, an analytical model including bending and torsion deformation for an anisotropic cylinder are derived. Physical forces controlling the bending and torsion deformation are carefully explored in Sec. 4.

## 2 The last stages of the *C. elegans* embryo just before hatching

*Caenorhabditis elegans* (*C. elegans*) provides an anatomically simple and integrated model to study the effect of physical forces (Brenner, 2003; Izquierdo and Beer, 2018; Palyanov et al., 2018). It has been reported that embryonic elongation in *C. elegans* occurs in two stages, *e*.*g*. 1.7-fold elongation due to epidermal actomyosin contractility in the first stage and 3-fold elongation induced by the coupling of epidermal actomyosin and muscle contraction in the second stage shown in Fig. 1(a) (Carvalho and Broday, 2020; Lardennois et al., 2019; Vuong-Brender et al., 2017). For the early stage, Ciarletta et al. (2009) provided a continuum model to explore the mechanical role of cellular filaments during embryogenesis and presented an axisymmetric elastic solution for the *C. elegans* embryo. Ben Amar et al. (2018) provided a model of active elongation material, combining pre-stress and passive stress in nonlinear elasticity, and predicted the contractile forces generated by the molecular motor myosin II for an elongation up to 1.7 times the initial length. They validated the analytical model by experimental measurements using laser fracture ablation, Vuong-Brender et al. (2017). The geometry of the deformation is simpler in this first stage since the cylindrical shape is perserved. This is explained by the ortho-radial position of the actin cables but the difficulty comes from the existence of different types of cells: the seam and the dorso-ventral cells, their different shear modulus and the fact that attached myosin molecules mainly concentrate in the seam cells. Lardennois et al. (2019) experimentally identified the kinases PAK-1 and SPC-1, which are crucial for actin remodelling in the second stage, and proposed that muscle contractions cause actin filaments to bend and sever. The remodelling regulated by SPC-1-PAK-1, with progressive shortening of actin filaments under the control of these factors, mediates a cellular viscoplastic process that promotes axial elongation. Based on the model proposed in Lardennois et al. (2019), Fang et al. (2021) established a unified dynamic model to explain how cellular anisotropy and plasticity, induced by actin filament alignment and severing/rebinding, dictate the elongation dynamics of *C. elegans* embryos.

**Figure 1.**
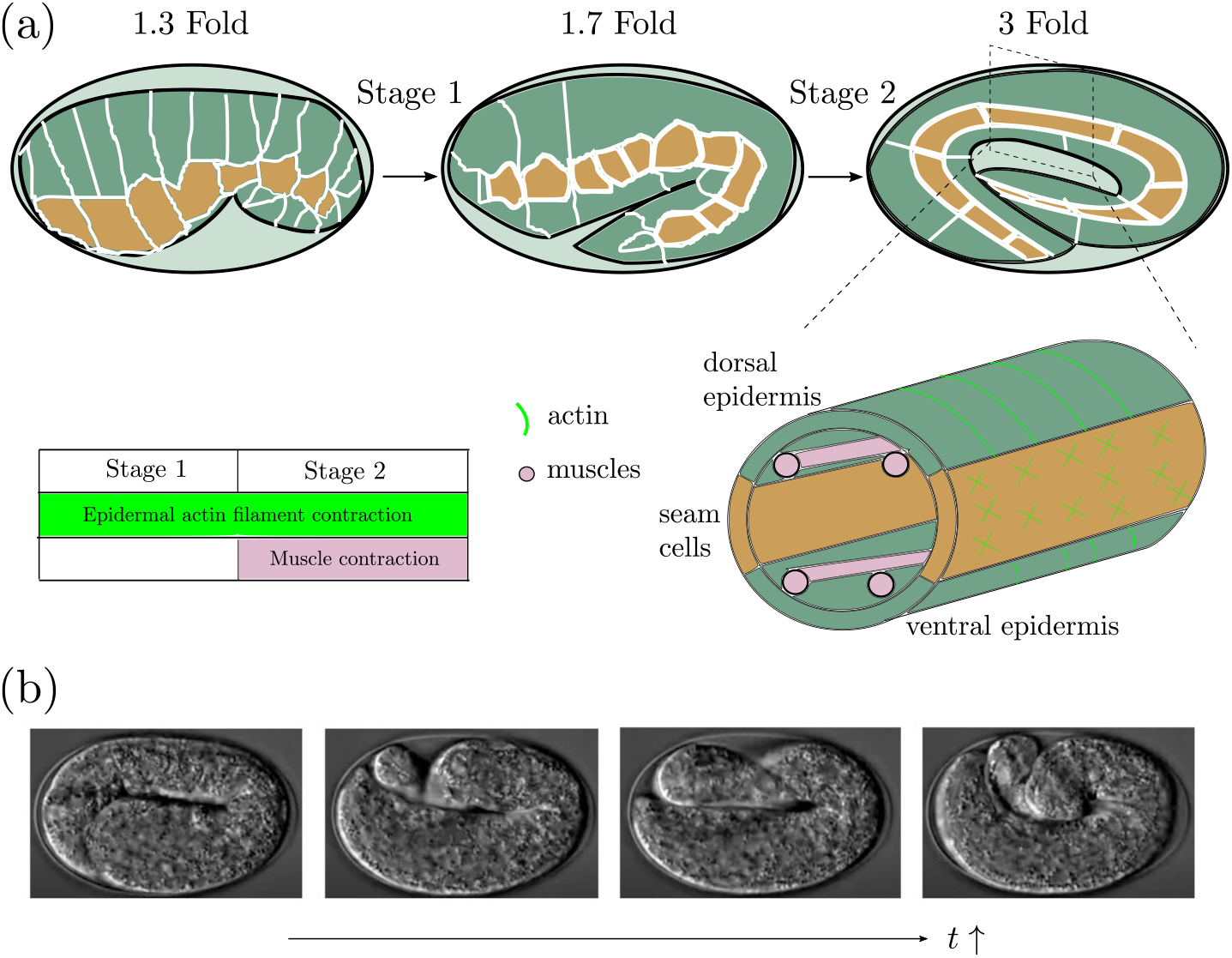
(a) *C. elegans* embryonic development is driven by epidermal actin filaments contractions during whole process and coupled with muscle contractions from 1.7-fold of initial length. (b) Obvious torsion can be observed during the evolutions of *C. elegans* embryonic development. (See https://www.youtube.com/watch?v=M2ApXHhYbaw)

Previous studies have demonstrated that the contractile force of myosin motors drives the first stage of elongation of the body axis of *C. elegans* analytically and experimentally (Ben Amar et al., 2018; Lardennois et al., 2019; Vuong-Brender et al., 2017). When the embryo has reached the second stage of elongation, the muscles underlying the epidermis become the main force generator as their contractile capacity starts around the twofold stage (Carvalho and Broday, 2020; Lardennois et al., 2019). In the *C. elegans* embryonic development at this stage, obviously complicated deformations including bending, rotation and torsion can be observed, as shown in Fig. 1(b). The rotations which limit the experimental investigations are due to the slight muscle deviation from the central axis (Dai and Ben Amar, 2023).

How to understand the complicated behaviour and predict the forces induced by the actin filaments or the muscles remains unclear. In the next section, we develop a novel analytical model that can predict the physical forces to effectively explain their complex nonlinear behaviours, which cannot be achieved by the existing models that focus on the cylindrical deformations under the passive stress (Goriely and Tabor, 2013; Sigaeva et al., 2018).

## 3 A mechanical model of bending and torsion of an anisotropic cylinder

Experimental dynamic observations show that *C. elegans* embryonic development involves rotational, bending and torsional deformations (as shown in Fig. 2) due to the epidermal actin filaments and to muscle contractions. These two networks of active filaments have different geometries and locations, independent of the mechanical energy they can develop. The experimental study by laser fracture ablation, which was possible in the first period of elongation (Ben Amar et al., 2018; Vuong-Brender et al., 2017) and which gives important results on the elastic stresses in the embryonic epidermis, becomes impossible once the muscles start their contractions. In fact, the embryo is twitching too fast and hinders fractures by laser ablation on the body in the second stage. It remains unclear how to estimate the forces induced at this stage. In this paper, we develop a mechanical model including bending and torsional deformations of an anisotropic cylinder to calculate the forces which occur in the second stage of the *C. elegans* elongation. Assuming the configuration shown in Fig. 3, the cross section includes four pieces to mimic the dorsal and ventral epidermal parts, and the muscles are considered at the interface to simplify the model. In fact, muscles are located quite at the border of the epidermis, slightly shifted in the vicinity of the DV cells with a tiny angular slope with the vertical axis. Here a morpho-elastic model is proposed for the evaluation of the stresses generated by the active filaments: actomyosin and muscles.

**Figure 2.**
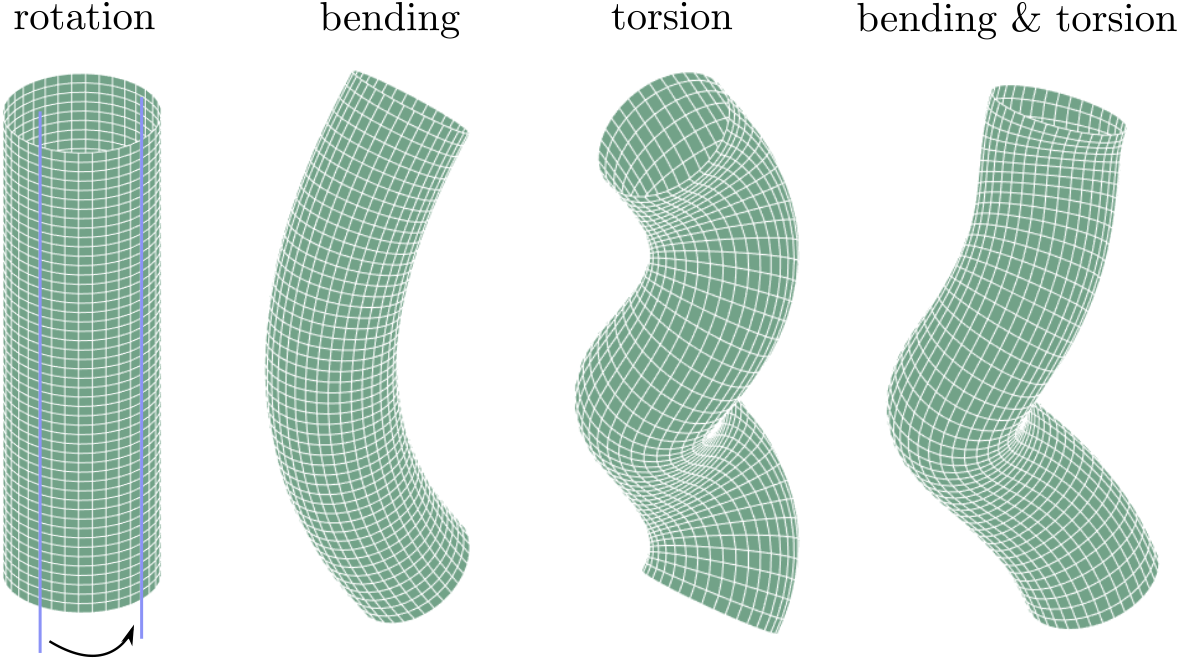
Schematic diagram of rotation, bending, torsion and bending plus torsion deformations for a cylindrical shell.

**Figure 3.**
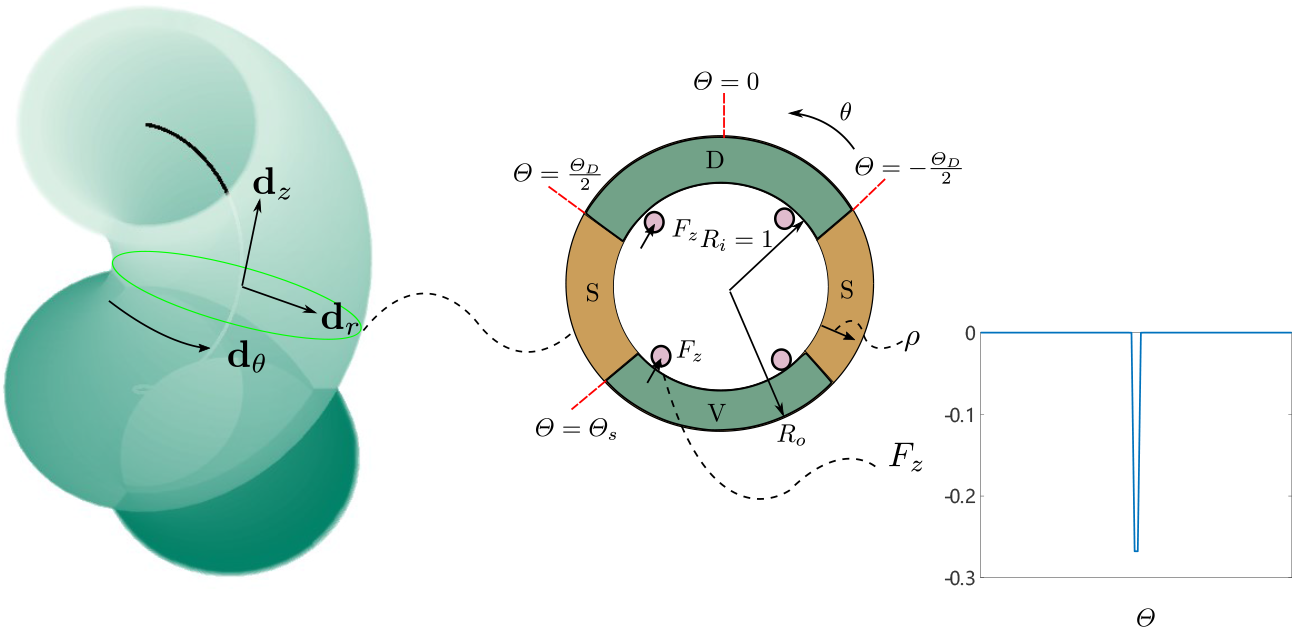
On the left, representation of the current configuration mimicking the deformation of the *C. elegans* embryo. In the middle, the cross-sectional geometry when the body remains cylindrical. Colors in the section represent different cell types: green for dorsal (D) and ventral (V) cells, yellow for seam (S) cells, and red for muscles. On the right, the localized active stress due to muscles near the boundary between S and DV cells.

### 3.1 Bending & torsion deformations

We consider the deformation from the initial straight tubular configuration to the current configuration, including bending and torsional deformations, as shown in Fig. 3. Noticing that the radius is smaller than the length even at the onset of embryonic elongation, we introduce a change in the scaling of the local cross section such that 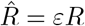, where *ε* is a small parameter of the order of 10% (Dai and Ben Amar, 2023). The mapping between the initial and the current configuration is then (Kaczmarski et al., 2022; Moulton et al., 2020)

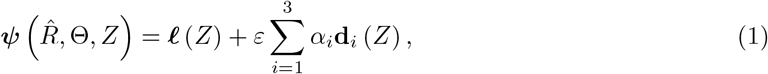

where α_*i*_ represents the deformation of the cross section, ℓ (*Z*) represents the centerline, and ℓ ^*′*^ (*Z*) = *ζ***d**_*z*_, *ζ* being the axial extension, and **d**_*i*_ indicates the local director basis, satisfying 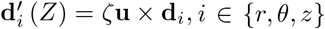. The Darboux curvature vector is defined as **u** = *u*_*r*_**d**_*r*_ + *u*_*θ*_**d**_*θ*_ + *u*_3_**d**_*z*_. In this way, we derive the deformation tensor in the local basis:

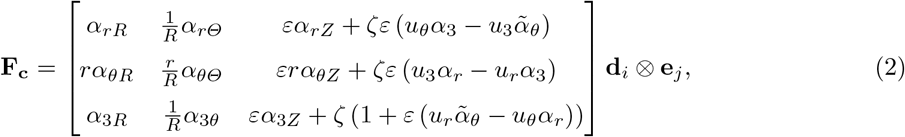

where the reference state is chosen at the end of the first stage, shown in Fig. 1(a), which shows an elongation reaching 1.7 and where α_*rR*_ = *∂*_*Rαr*_ and *j* ∈ {*R*, Θ, *Z*}. Of course, there is a pre-stretch tensor due to the first stage elongation, defined as **F**_**0**_ = Diag((*λ*_0_*λ*_0Z_)^*−*1^, *λ*_0_, *λ*_0Z_). This pre-stretch tensor takes into account both the ventral enclosure and the elongation due to actomyosin contraction. Then the elastic tensor **F** reads **F** = **F**_**c**_**F**_**0**_. In the local cross section, we approximate the deformation to the following form including enhancement/reduction and torsion for a sector of an annular wedge (Horgan and Saccomandi, 1999; Yavari and Goriely, 2021) as follows:

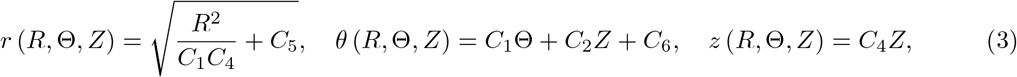

where *C*_*t*_ stands for the constants labelled with *t ∈* {1, 2, 4, 5, 6}. Finally, defining α_*r*_ = *r, α*_*θ*_ = *θ, α*_3_ = *h*(*Z*) and 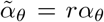 for a cylindrical geometry, the incompressibility constraint, Det[**F**] = 1, imposes *ζ* = *C*_4_ at the leading order in *ε*. So *ζ* is a constant.

We choose a general constitutive relationship for the energy density valid for fibrous materials (Horgan and Saccomandi, 2005; Horgan, 2015; Merodio et al., 2007).

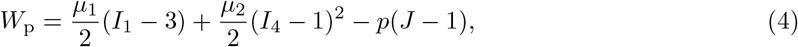

where *µ*_1_, *µ*_2_ are the shear moduli, the elastic volume change is defined by*J* = Det[**F**], and the parameter *p* is a Lagrange multiplier, which is determined by the boundary condition 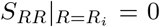, where *R*_*i*_, is the inner radius. The first strain invariant *I*_1_ is given by *I*_1_ = tr**C**, where **C** = **F**^*T*^ **F**, and the anisotropic invariant satisfies *I*_4_ = **a**_0_ *·* **Ca**_0_, where **a**_0_ is the unit vector indicating the direction of the fibre reinforcement in the reference configuration. The passive nominal stress **S**^*p*^ can be expressed as:

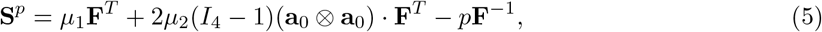

then, the passive Cauchy stress *σ*^*p*^ is given as:

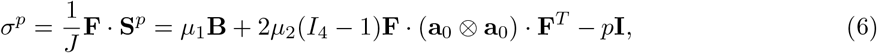

where **B** = **FF**^*T*^, and **I** is the (3 × 3) identity tensor. Since the actin filaments are distributed in average along the circumferential direction, the unit vector **a**_0_ is represented by **a**_0_ = {0, 1, 0} and the passive stress can be obtained from Eq.(6). Our formulation of active matter consists of an active stress in opposition of an active strain (Dai and Ben Amar, 2023), and the Cauchy stress becomes the superposition of a passive and of an active stress including an isotropic and a deviatoric tensor 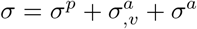 (Ben Amar et al., 2018; Prost et al., 2015). The deviatoric part of the active stress *σ*^*a*^ has the following form:

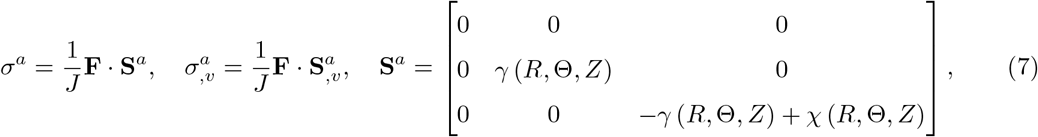

where *γ* (*R*, Θ, *Z*) is one component of the active stress induced by the actomyosin motors and *χ* (*R*, Θ, *Z*) represents the local active stress due to the muscles, in our case *J* = 1. The active stress *σ*^*a*^ is postulated phenomenologically to represent the effect of local contractions (Goriely, 2018), since only **S**^*a*^ is intrinsically related to the initial geometry of the active components. To mimic the localization, we choose for the muscle representation: 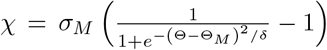, which is plotted in Fig. 3, where *δ* is a small parameter, taking 0.001, and Θ_*M*_ represents the angular position of the muscle. The muscles are located in the dorsal and ventral parts, near the interfaces with the seam parts, with Θ_*M*_ taking values close to Θ_*D*_*/*2 or Θ_*S*_ on the left, and *−*Θ_*D*_*/*2 or *−*Θ_*S*_ on the right as shown in Fig. 3. The volumetric part 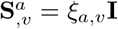 can be included into the Lagrange multiplier *p*. Without the presence of body forces, the equilibrium equation can be expressed as:

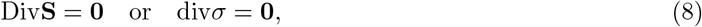

where Div(·) and div(·) are the divergence operators in the initial and current configurations, respectively. Corresponding boundary conditions at the free edges for stress satisfy

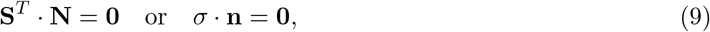

where **N** and **n** are normal unit vectors in the initial and current configurations. The equilibrium equations are given by div*σ* = **0** and it reads (Ogden, 1997):

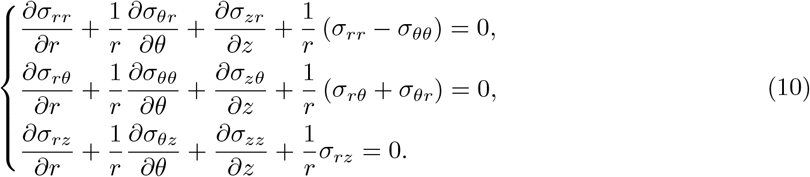

The above equation remains valid even if the Cauchy stress is not symmetric, so in the case of active forces generating volumetric torque.

### 3.2 Geometry of the embryo filament networks and constraints on parameters

The epidermis of the embryo has two types of cells: the seam cells and the dorso-ventral cells. The DV cells are stiffer, the fibres are more ordered, but they do not seem to have molecular motors attached to the fibres. So *σ*^*a*^ *∼* 0 in the DV cells, unlike the seam cells. As for the muscles attached close to the inner surface of the epidermis, they are roughly parallel to the vertical axis and are located near the boundary between the seam and DV cells (see Fig. 3). Here we will represent the compressive activity of these muscles by an active stress acting on the interface between the seam and the DV cells. We define the midpoint of the dorsal region as the coordinate origin. Note that the muscles are not always active and that they alternately work on one side, then relax and work on the other side. Here we consider that all the morphogenetic processes take place in the epidermis, but it is surrounded by a stiff and extremely thin cuticle and, in addition, the epidermis surrounds a ventral zone containing organs such as the intestine.

Expanding all the variables *V* (*V* = *σ, p, γ, χ*, …) in the small parameters *ε*, one has

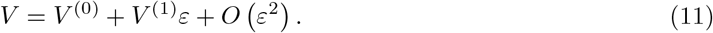

To simplify the analysis, we also expand each variable *V* ^(*i*)^ with {*i, j*} *∈* {0, 1} at each order along the thickness *R* = 1 + *ρ*, where *ρ* is a small variable, then one has

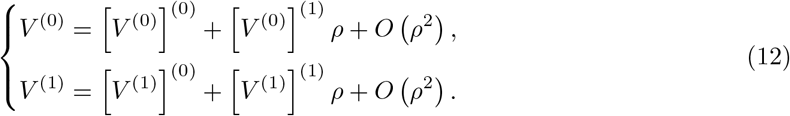

Since, at each order, the amplitude of muscle stress gradually decreases along the radial direction, we assume:

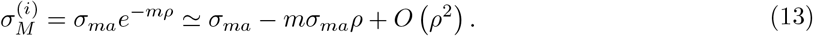

The ortho-radial stress component *S*_ΘΘ_ and the ortho-vertical stress component *S*_Θ*Z*_ are continuous accross the frontier between the S and DV cells. The continuity conditions read

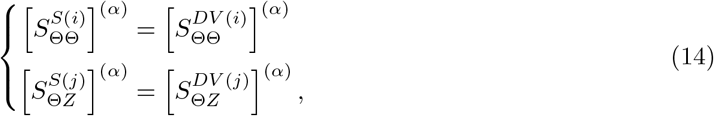

with {α, *β*} ∈ {0, 1}. Then, the coefficients of the active stress can be determined if the constants *C*_*t*_ are known. The active stress induced by molecular motors only works on the seam parts and is zero in the DV parts, while the active stress induced by muscle activity only works on the DV parts instead of the seam parts. As the molecular motors contract the seam parts, the angular stretch 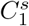 will gradually decrease and become less than 1, whereas the angular stretch 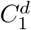 for the DV parts will gradually increase and become greater than 1. Since the experimental data on the circumferential length in the last stage are limited, and since 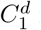 follows the same tendency as in the first stage, we adopt a similar variation of 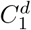 with *C*_4_ as in Ben Amar et al. (2018). To close the cylinder, the following geometrical constraints must be satisfied:

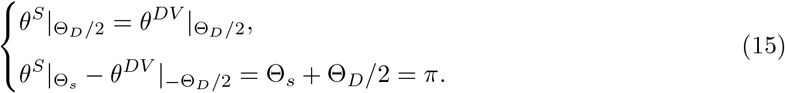

Taking 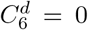, the constants 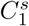 and 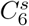 can be determined. The tendency of 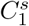 and 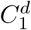 as the elongation proceeds are shown in Fig. 4. Remember that *C*_4_ is the elongation after the first stage and the total elongation is in fact: 1.7*C*_4_ after the ventral enclosure.

**Figure 4.**
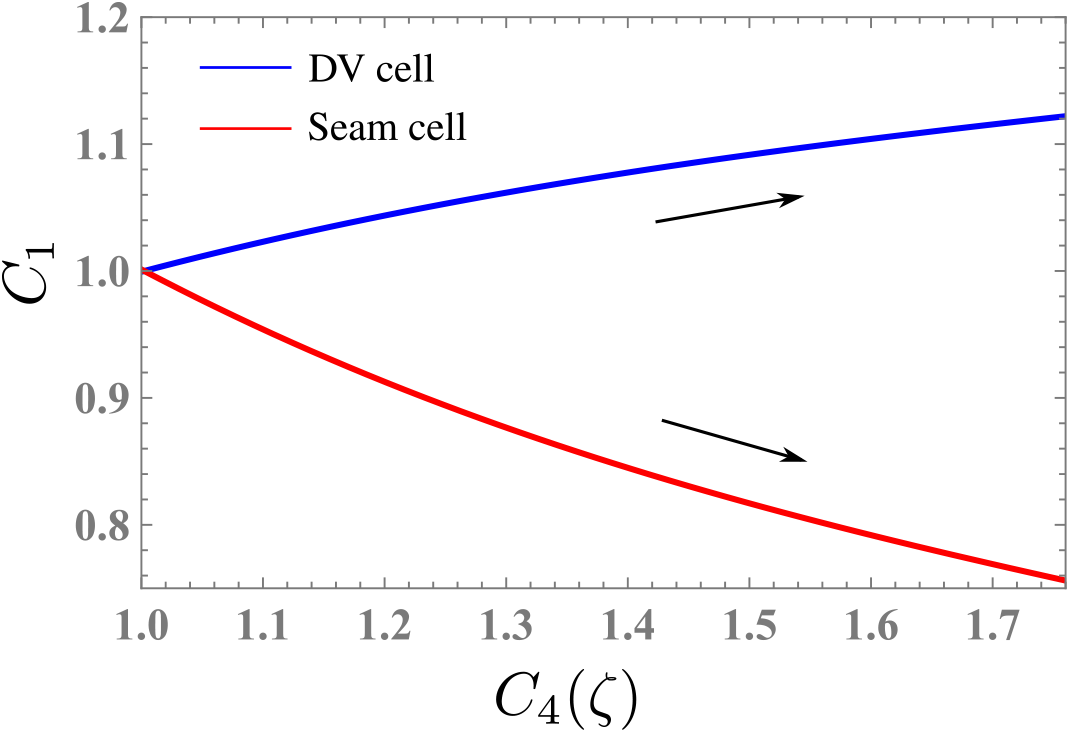
The angular stretch *C*_1_ of the seam and DV domains which varies with the elongation represented by 1.7*C*_4_, since the prestretch is *λ*_0*Z*_ = 1.7 at the beginning of the second stage.

Imposing that the inner interface for *R* = 1 remains perfectly circular gives a relation between the coefficients introduced in Eq.(3) that reads

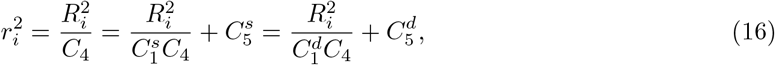

that is

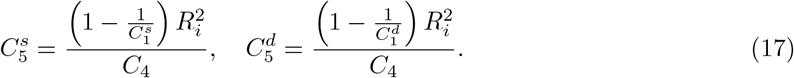

## 4 Results

### 4.1 Active stress generating bending deformation

We start with a simple example capable of generative bending deformation due to active stresses but without torsion as shown as Figs. 5 (b) and (e). The state of strain, when it reaches 1.7 times, is chosen as the reference configuration, and it marks the beginning of muscular activity and more complex patterns of deformation. The bending deformation will be induced with *u*_1_ = 0, *u*_2_ = constant, u_3_ = 0 and one will obtain *u*_*r*_ and *u*_*θ*_ based on the conversion relationship between cylindrical and rectangular coordinates in Eq. (31) of the Appendix B. Assuming that the deviatoric part of active stresses is *γ*^(*i*)^ = *a*_*i*_ *R* + *b*_*i*_ and 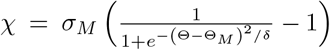 with 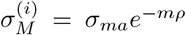, then, based on the first equilibrium equation in Eq. (10) and boundary condition *S*_*RR*_|_*R*=1_ = 0, one can obtain the Lagrange multiplier *p*: *p*[*R*, Θ]^(*i*)^ at each order. As yet mentioned, the active stress *γ*^(*i*)^ only works on the seam parts. The detail derivations for Lagrange multiplier *p* are presented in Appendix A.

**Figure 5.**
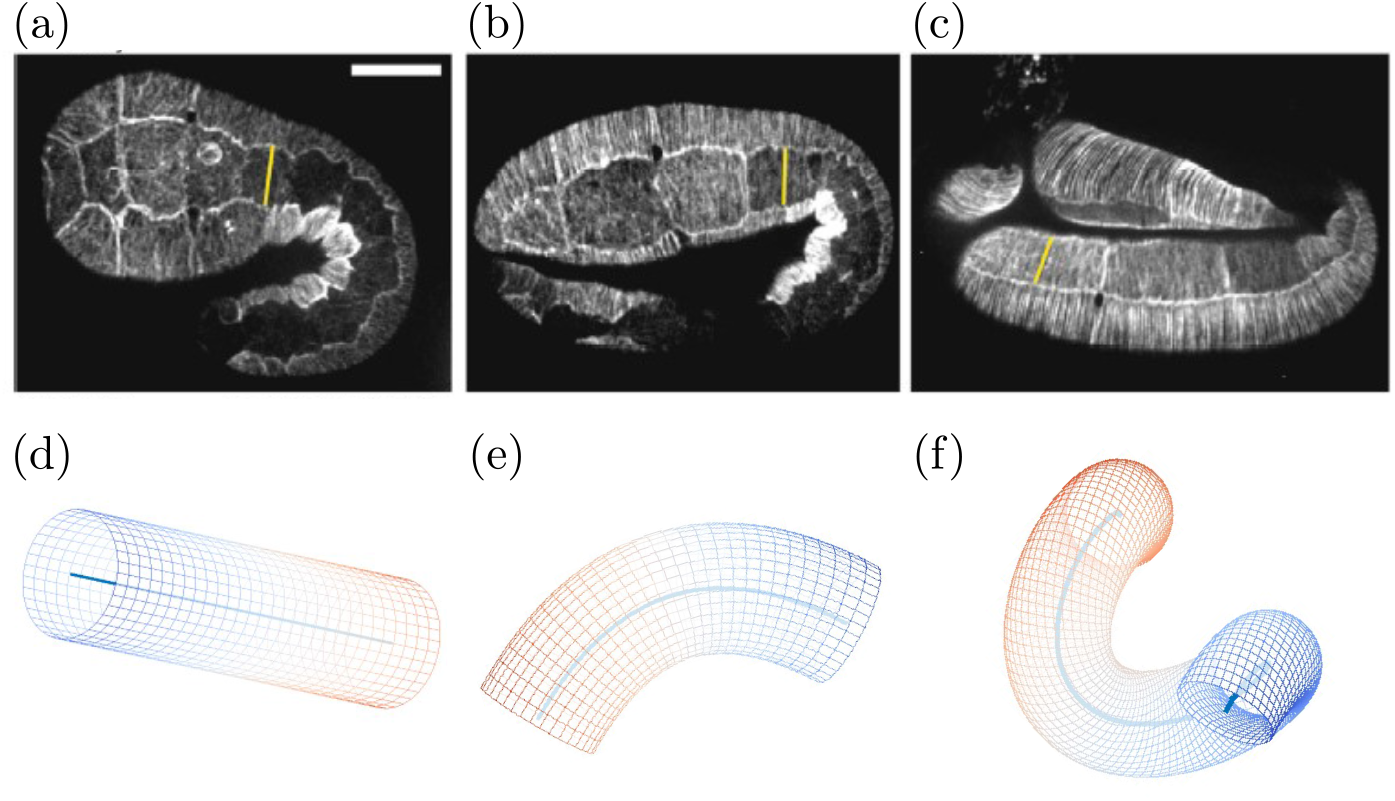
(a)-(c) Three elongation stages corresponding to 1.7-fold, 2-fold and 3-fold (Lardennois et al., 2019). (a) and (d) corresponds to the beginning of stage 2. (d)-(f) simplified configurations of the body shape corresponding to (a) to (c).

The second and third equilibrium equations (10) at leading order and first order of *ε* are satisfied automatically with the condition *C*_2_ = 0 and *u*_3_ = 0 for bending deformation and by α_3_ = 0.

Next, the active stresses *σ*^*a*^ and 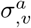 are determined by the continuity conditions and taking 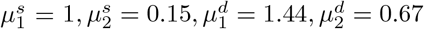 (Ben Amar et al., 2018). One has

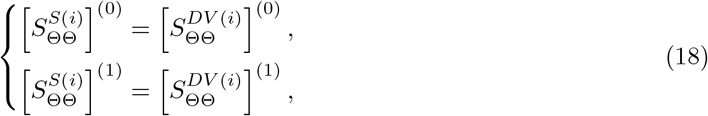

then, *a*_*i*_ and *b*_*i*_, the two constants of the active components *γ*^(*i*)^ are obtained, which have the form *f* (*ζ*). The deviatoric part of active stress *γ*^(*i*)^ = *a*_*i*_*R* + *b*_*i*_ induced by the actomyosins can be achieved. The corresponding isotropic part 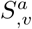 has the following form

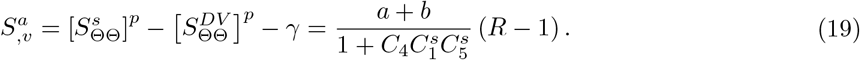

The active stress induced by actomyosin molecular motors demonstrates varying patterns in response to changes in elongation and radius, as illustrated in Fig. 6. Notably, the deviatoric component of the active stress, denoted as *γ*, undergoes increase in the process. Importantly, an increase in elongation results in an augmentation of the deviatoric active stress *γ*, as expected and as observed in Fig. 6(a). In Fig. 6(b), the volumetric component of the active stress, denoted as 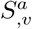, exhibits similar tendency, with its magnitude increasing concurrently with elongation at *R* = 1.25. The compressive nature of the passive stress observed in the seam and dorsal parts, presented in Fig. 6(c), is also noteworthy. Furthermore, the overall active stress, comprising both isotropic and deviatoric components, as depicted in Fig. 6(d), displays a increasing trend with increasing elongation and radius. This comprehensive analysis contributes valuable insights into the intricate interdependence of active stress with elongation and radii.

**Figure 6.**
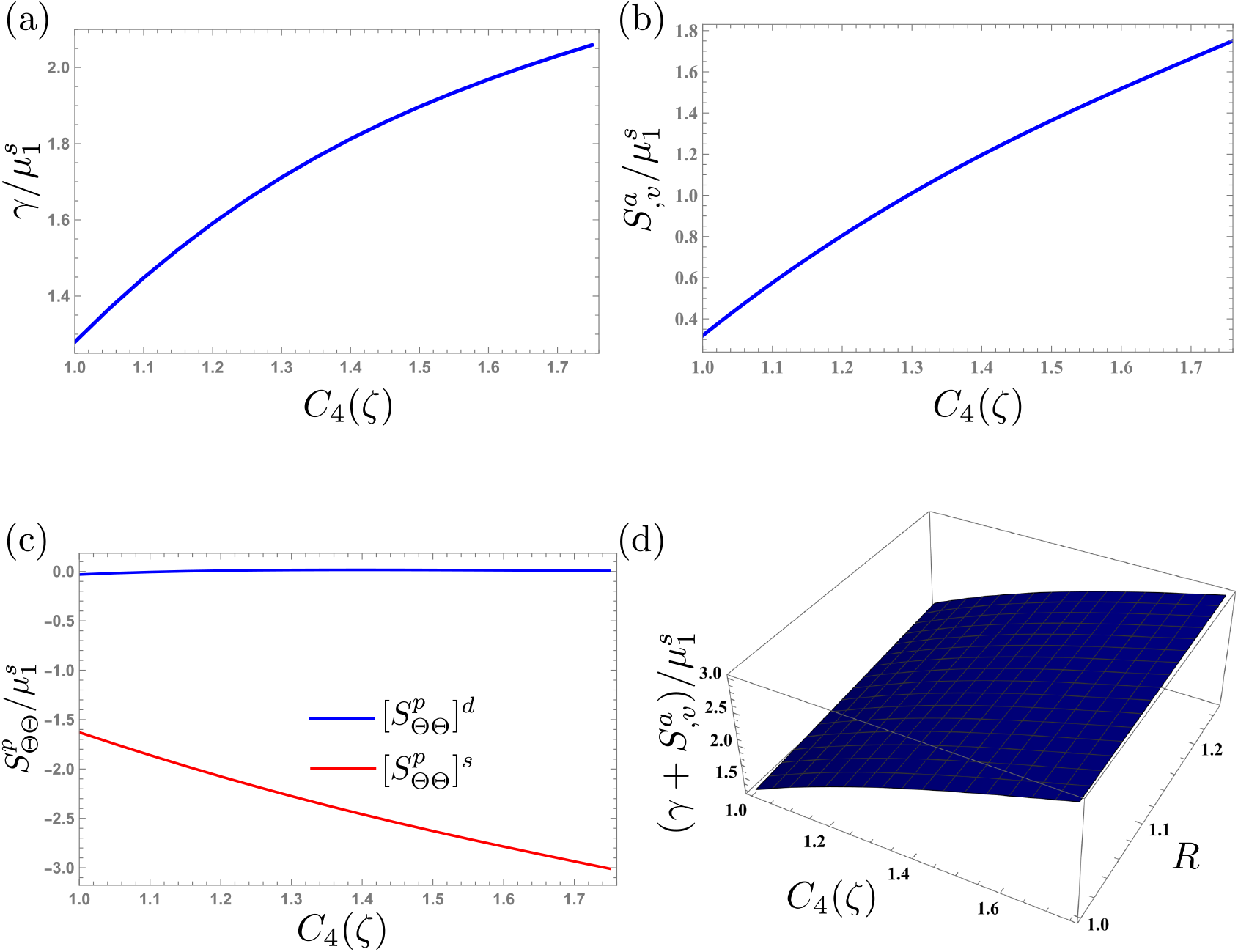
(a) The deviatoric component of the actomyosin active stress *γ* varies with elongation at *R* = 1.25. Obviously, as the elongation increases, the deviatoric component also increases. (b) The isotropic component of the active stress, denoted as 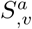, exhibits an increase with increasing elongation at *R* = 1.25. (c) The passive stress 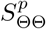 of the seam and dorsal parts at *R* = 1.25. (d) The active stress varies with the elongation and radius. From the outer surface to the inner surface, the active stress induced by the actomyosin network decreases, which is consistent with the geometry of the cables. Prestretch parameters are: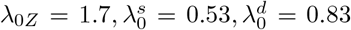. Note that the reference configuration is determined by the first elongation that reaches 1.7 times (*λ*_0*Z*_ = 1.7), and indicates the beginning of muscle engagement.

The active stress χ ^(*i*)^ induced by muscles can be determined by the free boundary condition on the top of the interface

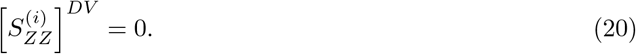

We subsequently investigate the impact of elongation, bending curvature and radius on the active stress induced by muscular activity, as illustrated in Fig. 7. The results reveal that an increase in elongation leads to an augmentation of the magnitude of the active stress due to muscles, denoted as χ (see Fig. 7(a)). Additionally, the active stress χ exhibits compressive characteristics, demonstrating an augmentation in magnitude concurrent with the increase in bending curvature *κ* at *R* = 1. In Fig. 7(b), the magnitude of the active stress χ gradually decreases from the inner surface to the outer surface.

**Figure 7.**
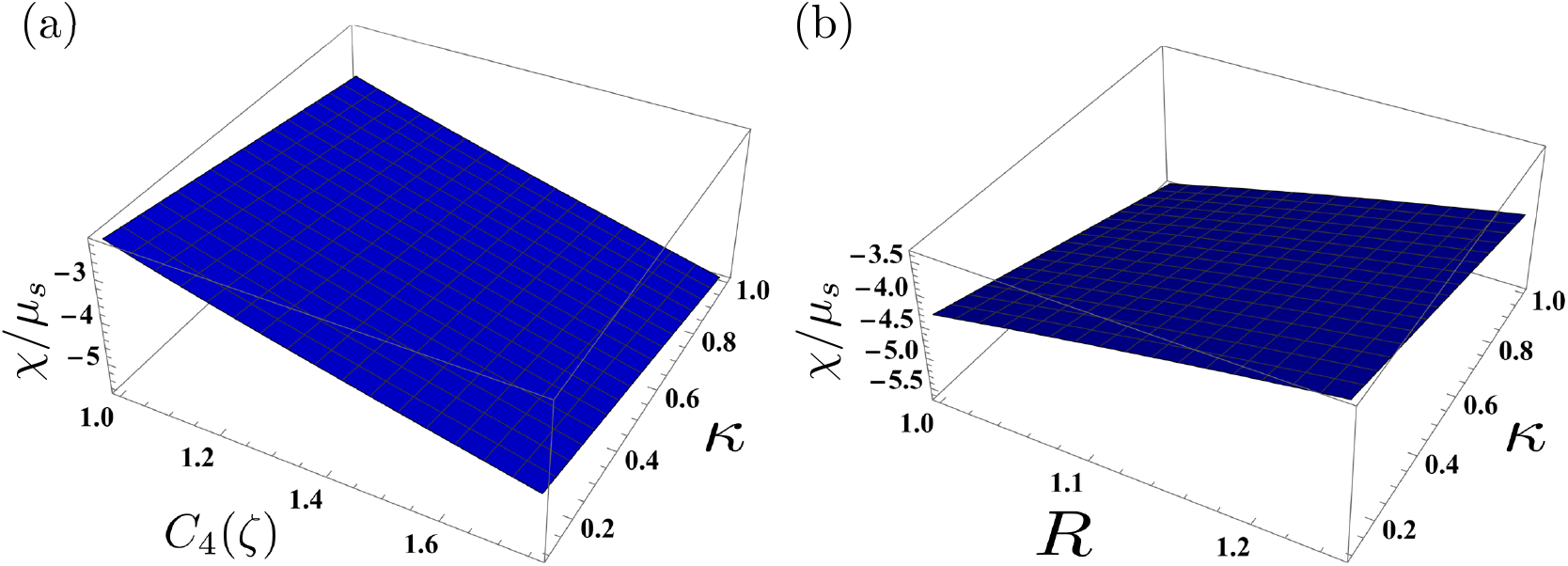
(a) The active stress induced by muscles undergoes variations in response to elongation and bending curvature at *R* = 1, showing that the magnitude of the active stress increases with the increase of elongation and curvature. (b) The active stress induced by muscles varies with radius and bending curvature at *C*_4_ = 1.76. The magnitude of the active stress due to muscles is smaller from the inner surface to the outer. A significant value of *ε* = 0.3 is chosen to amplify the effect of bending curvature. Prestretch parameters: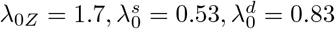.

Finally, to illustrate that the different sections still maintain closure after deformations so that the embryo integrity is maintained, we provide a schematic diagram for the cylinder with an elongation *C*_4_ = 1.2 and a bending deformation *κ* = 0.1. When the curvature and arc length of the bent cylinder are known, the centerline can be determined. The results demonstrate that our calculations satisfy the geometric constraints and that effectively all parts of the cylindrical shell remain connected.

### 4.2 Physical force generating extrinsic torsion

In the scenario shown in Fig. 2, the cylindrical structure undergoes a significant torsional deformation along its axis. This torsional deformation can be characterized by the curvature: *u*_*r*_[*Z*], *u*_*θ*_[*Z*] and *u*_3_. Note that the torsion of the embryo is observed for a significant elongation in a confined space. The geometry of the active stresses studied previously cannot simply explain a torsion that requires a moment applied at the top of the shell: this moment can be realized by circular muscles acting at the neck of the embryo. Assuming that they are much more active in the late elongation, we neglect the active network operating before and consider only this moment. We then carry out our calculations under these conditions in order to understand the torsional response of the system. The moment *M* is defined as (Hamdaoui et al., 2014)

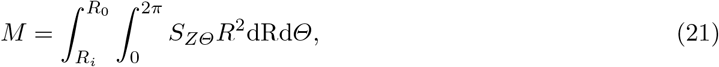

where the shear stress *S*_*ZΘ*_ has the following form which can be deduced from Eq. (5):

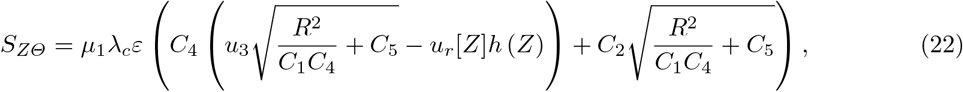

where *C*_2_ = *C*_4_ τ and τ represents the torsional deformation per unit deformed length. Note that in the DV and seam parts the shear stresses are different. The second equilibrium equation (10) at the first order of *ε* yields *h*[*Z*] = *−u*_*r*_[*Z*]. To facilitate the analysis, we consider a specialized centerline characterized by a helical curve with *u*_3_ = 0 and τ = *ω* (Moulton et al., 2020). The detailed derivation correlating the Darboux curvature **u** with the curvature *κ* and torsion *τ* of the centerline is provided in Appendix B. Consequently, *u*_*r*_[*Z*] = *κ* sin (τ *Z*).

The variations in moment magnitudes *M* with elongation and torsional deformation are depicted in Fig. 9. Evidently, an escalation in elongation leads to a corresponding increase in moment magnitude.

**Figure 8.**
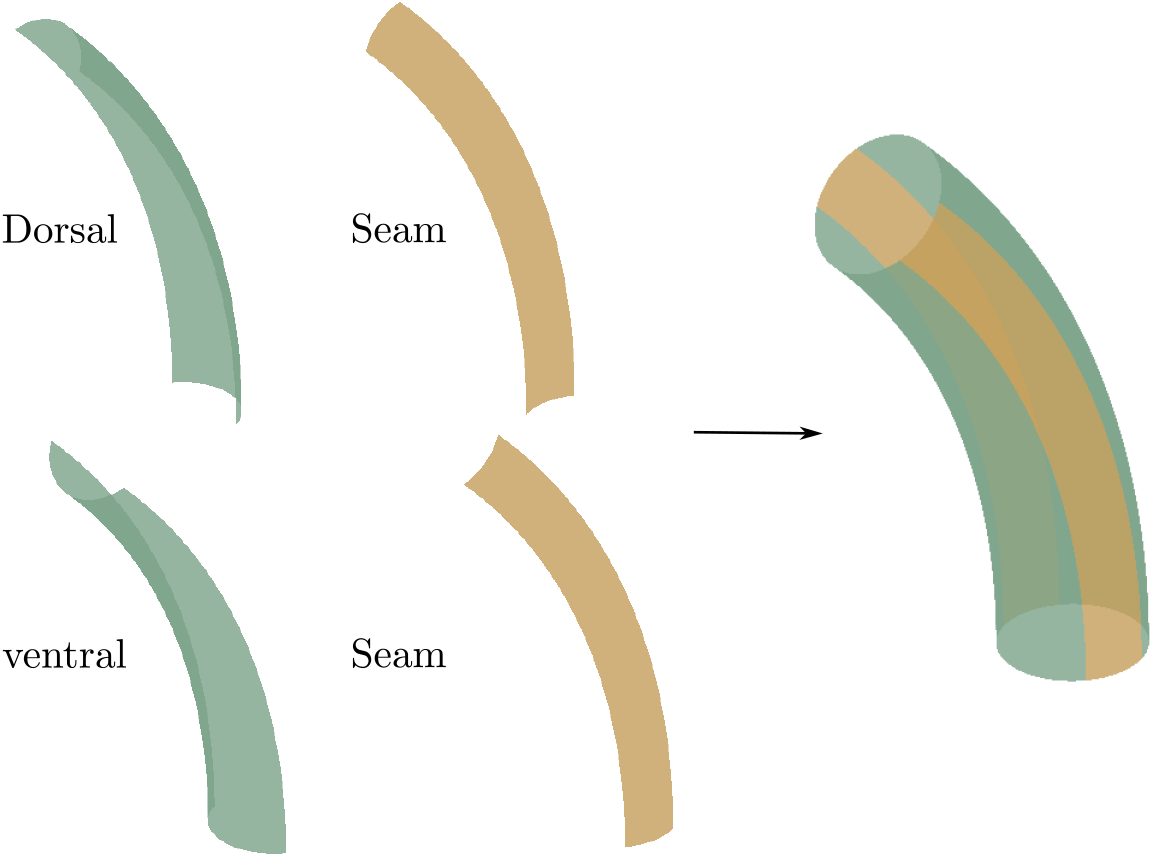
The demonstration depicting four segments closing around the bent cylinder validates the fulfillment of geometric constraints through our calculations. Deformation parameters: *C*_4_ = 1.2 and *κ* = 0.1.

**Figure 9.**
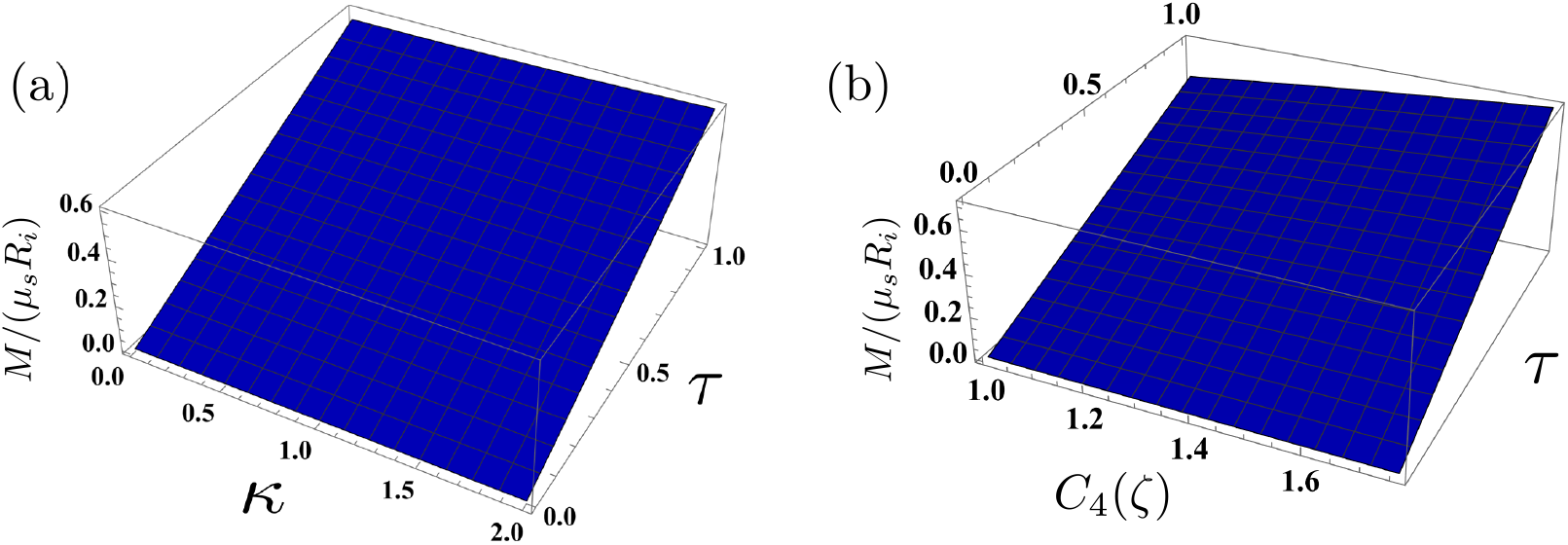
Variation of *M*, the moment at the top of the body shape, with both elongation and torsional deformation, demonstrating a positive correlation where moment magnitudes escalate as elongation and torsion increase. (a) Given *C*_4_ = 1.2, the moment *M* escalates proportionally with the augmentation of curvature and torsion. (b) When *κ* = 1.35, the moment *M* amplifies in correlation with elongation and torsion, while maintaining *ε* = 0.3. Prestretch parameters: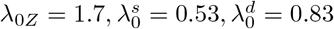.

In addition, the increase of torsional deformation also contributes to increased moment levels. To demonstrate the continued closure of various sections despite deformations, thus ensuring the integrity of the embryo, we present a schematic diagram in Fig. 10 depicting a cylinder elongated to *C*_4_ = 1.2 and subjected to torsional deformation of *C*_2_ = 0.96, with curvature *κ* = 1.35 and τ = 0.8. Our results validate that our computations adhere to geometric constraints, ensuring the connection of all segments within the torsional cylindrical shell. However, this analysis restricts on the torsion of the epidermis maintaining the axis of symmetry and the intrinsic torsion is discarded.

**Figure 10.**
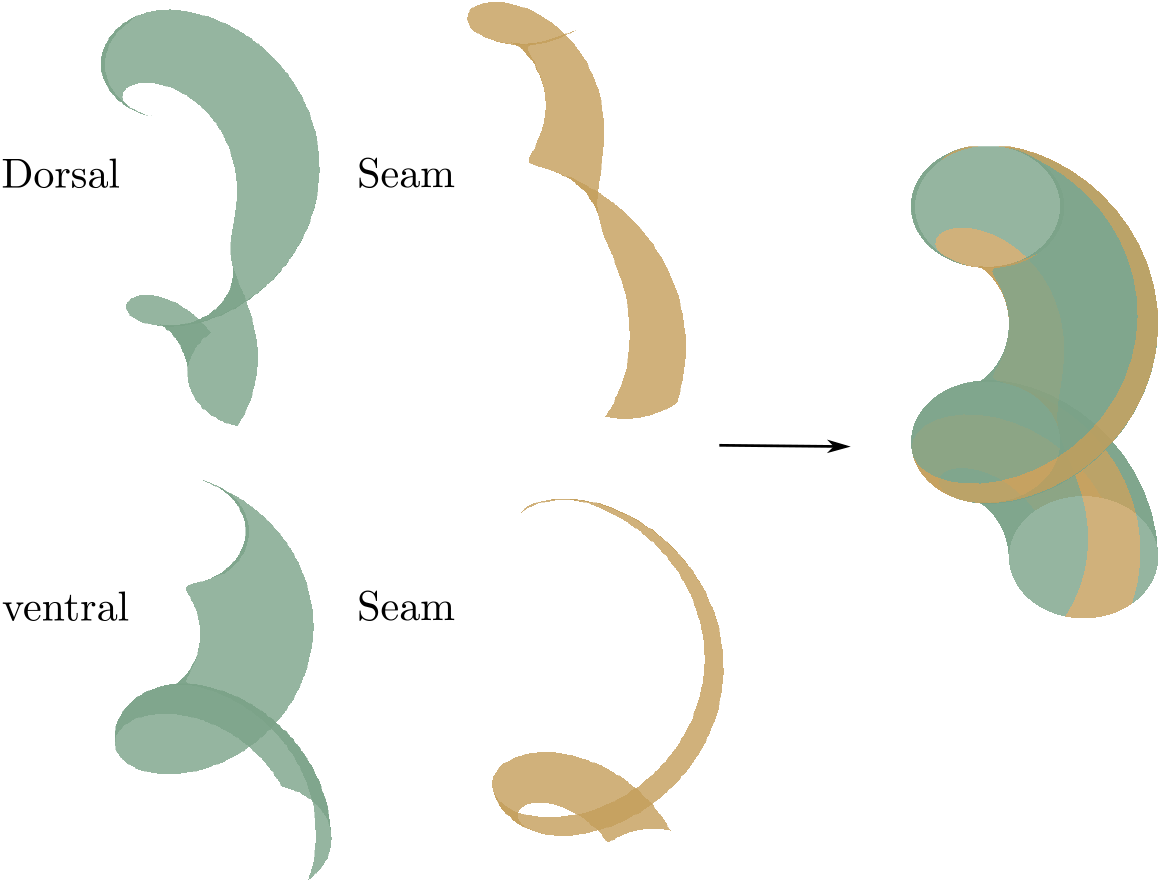
The demonstration depicting four segments closing around the torsional cylinder validates the fulfillment of geometric constraints through our calculations. Deformation parameters: *C*_4_ = 1.2 and *C*_2_ = 0.96, with curvature *κ* = 1.35 and τ = 0.8. Function of the centerline: **ℓ** (*Z*) = (0.6 sin (1.5*Z*), *−*0.6 cos (1.5*Z*), 1.2*Z*). The detail calculation for corresponding Darboux curvature **u** are presented in Appendix B

## 5 Conclusion

Here, we have uncovered the impact of epidermal actomyosin and muscle contraction on the extension of the body axis in the midst bending and torsional deformations during embryonic morphogenesis, in the absence of cellular rearrangement and proliferation. Using a mechanical model of bending and torsion for an inhomogeneous cylinder, we evaluated the active stress induced by actomyosin and muscle contractions. Our results reveal that the active stress induced by actomyosin molecular motors or muscles increases with elongation and bending curvature, while also varying with radius. In addition, both elongation and torsional deformation contribute to increased moment magnitudes. These results highlight the complex interplay of biomechanical factors in modulating embryonic deformation. In this article, due to the complexity of the embryo geometry, we separate the first stage of elongation due to the actomyosin network from the second stage of bending and torsion where actomyosin superimposes on lateral muscles. We mimic the deformations of the first stage by a pre-strain that we incorporate into the study of the second stage, where the active networks are then represented by active stresses. All together, the formalism of finite elasticity with pre-strain, preferred here to pre-stress, and active stresses, is able to quantify the strong deformation of the *C-elegans* embryo before hatching.

## Acknowledgements

This paper is dedicated with warm wishes to our friend Peppe Saccomandi on the occasion of his 60th birthday. Peppe is an extremely brilliant and talented mechanician and biomechanician and we learn a lot from his conference talks and articles. The authors acknowledge the support of the ANR (Agence Nationale de la Recherche) under the contract EpiMorph (ANR-2018-CE13-0008). They also acknowledge discussions with Alain Goriely, Kelly Molnar and Michel Labouesse.

## A Supplementary of the derivation

The deformation gradient **F** is expressed as

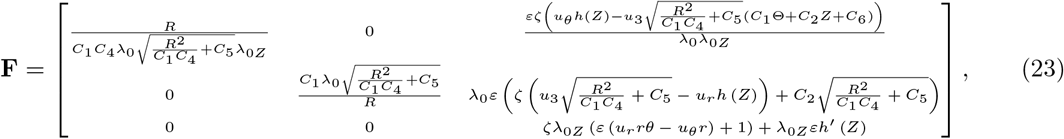

then, the determinant of deformation gradient **F** is expressed as the series of *ε*

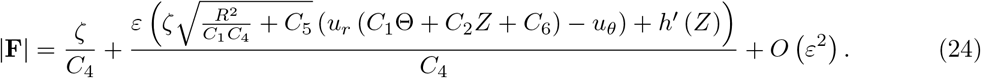

Assuming incompressibility, Det[**F**] = 1 and for simplification, only the zero order is considered: *ζ* = *C*_4_ to have Det[**F**] ≈ 1. Based on the Eqs. (2) and (5), one can calculate the passive stress as following

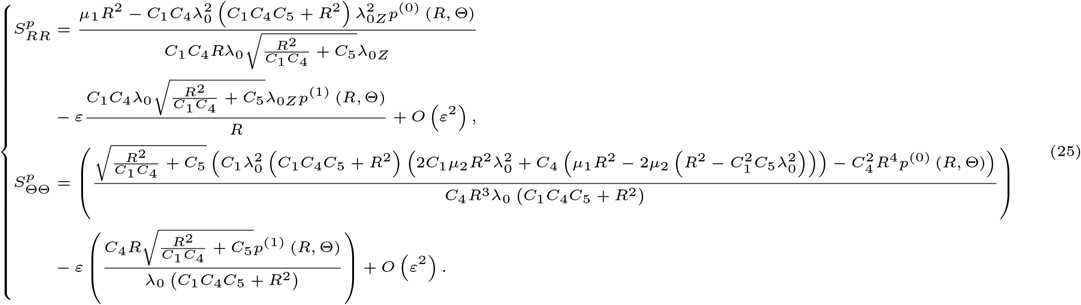

Similarly, the passive Cauchy stress can be obtained. Then, the stress including active stress can be achieved from 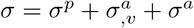. Next, we consider equilibrium equations at the leading order and the first order of *ε* separately.

For the leading order of *ε*, from the first equation of equilibrium equations (10), one has

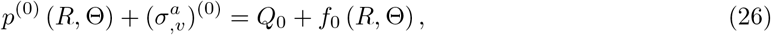

where *Q*_0_ is a parameter determined by boundary condition 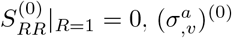 is the part including active stress coefficients *a*_0_, *b*_0_, and *p*^(0)^ can be separated by giving *a*_0_ = 0, *b*_0_ = 0.

Similarly, for the first order *ε*, from the first equation of equilibrium equations (10), one has

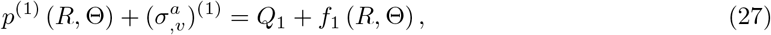

where *Q*_1_ is a parameter determined by boundary condition 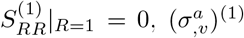 and *p*^(1)^ can be determined by the same way as the zero order.

## B Curvature and torsion of the centerline

For the given deformation of the centerline

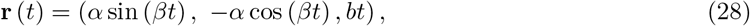

the curvature *κ* and torsion *τ* of the centerline can be calculated as

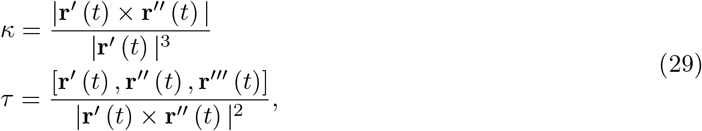

then, one has

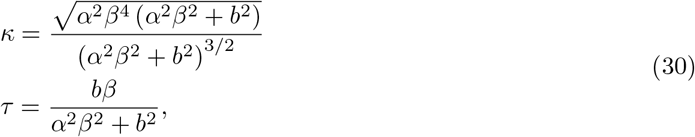

then, one will has 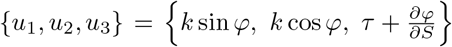, where *∂φ/∂S* represents the rotation of the local basis with respect to the Frenet frame as the arc length increases.

Conversion relationship between cylindrical coordinates and rectangular coordinates has

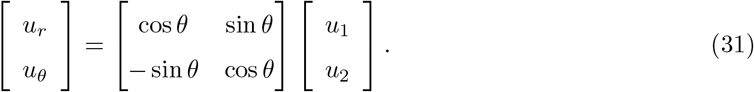

